# Lessons and recommendations from three decades as an NSF REU Site: A call for systems-based assessment

**DOI:** 10.1101/162289

**Authors:** Andrew L. McDevitt, Manisha V. Patel, Aaron M. Ellison

## Abstract

NSF’s Research Experiences for Undergraduates (REU) program supports thousands of undergraduate researchers annually. REU sites operate independently with regards to their research mission and structure, leading to a complex educational milieu distinct from traditional classrooms and labs. Overall, REU sites are perceived as providing highly formative experiences for developing researchers. However, given improved assessment practices over REU’s three decades, best practices for student learning and evaluation of long-term impacts remain limited. To address this limitation, we recommend the use of systems-based theoretical frameworks when studying REU programs. We outline how one such framework, cultural-historical activity theory (CHAT), could inform the collection of assessment data. Among other strengths, CHAT guides collection of quantitative and qualitative information that can help characterize REU programs in an educationally meaningful context. Adoption of CHAT and similar approaches by REU Sites could improve dialogue among programs, encourage collaborations, and improve evidence-based practices.

## Introduction

Undergraduate research experiences that provide authentic opportunities in research laboratories and field sites are a widely accepted way to strengthen student preparation within scientific disciplines (Kuh 2008). One of the first programs to support these experiences in the United States was the Undergraduate Research Participation Program, which the National Science Foundation (NSF) began in 1958 and eliminated in 1982 because of educational budget cuts (Neckers 1982). In 1987, NSF initiated the Research Experiences for Undergraduates (REU) program following the paid internship model of the Undergraduate Research Participation Program. Since then, REU has been one of the largest supporters of undergraduate research programs; $1.12 billion was invested in supporting thousands of undergraduates each year between 2002 and 2017 through both REU Site and REU Supplement awards (Figure S1).

Individual REU sites are defined by their intellectual themes and the community of researchers that they bring together. Although a common goal of the REU program may be to prepare undergraduate students for careers in STEM fields by providing scientific and engineering research opportunities, the design of these educational experiences depends on the goals and values of each site. Sites vary in their personnel, infrastructure, intellectual pursuits, and the student populations they serve. Sites also vary in how well they evaluate whether they are achieving their goals.

At least through 2010, individual REU sites selected and managed their own assessment protocols. Individual site assessments were unique case studies(Medina-Borja et al.) based on results of internally developed surveys (McDevitt et al. 2016) and participant surveys (Seymour et al. 2004). Qualitative data from these surveys elicited insights about student experiences but the data were neither representative nor a random sample and were expensive to collect. Quantitative surveys create less of a burden on programs and were widely used (Linn et al. 2015), but the surveys often consisted of conceptually ambiguous questions and, because they were developed by individual programs for internal use, were rarely validated, and are difficult to compare with other programs (Linn et al. 2015, McDevitt et al. 2016).

NSF also supported the development of assessment tools and examination of assessment data across REU Sites that have been used to evaluate their effectiveness. For example, by the early 2000s, it was clear that undergraduate research programs successfully recruited women (Kardash 2000, Liang et al. 2002) and minority students into STEM fields (Foertsch et al. 2000, Gregerman et al. 1998, Nnadozie et al. 2001). In 2003, NSF aligned REU program goals with these new findings and outlined REU funding priorities in their yearly congressional budgets (Figure S1A). The passage of the America COMPETES Act of 2010 (P.L. 111-478 §514) strengthened initiatives to reach diverse participants, especially from institutions where STEM research opportunities are limited. It also mandated the tracking STEM matriculation and employment of participants for at least three years following graduation. Around the same time, the Biology REU Leadership Committee began using a common assessment tool, the Undergraduate Research Student Self-Assessment (URSSA; Hunter et al. 2009), to evaluate common goals and improve communication about BIO-REU programs (BIO REU Leadership Committee 2010).

The flexibility afforded to REU sites by NSF encourages innovative pedagogical approaches but also increases the heterogeneity across programs. In contrast, surveys such as URSSA were developed with specific programmatic goals in mind. Both individual site-based surveys and cross-site surveys like URSSA serve their intended purpose, but the lack of theoretical underpinnings limits our ability to relate our findings to the broader literature on education or to compare them more meaningfully to data from other REU Sites. We experienced this limitation when examining 10 years of pre/post surveys from the Harvard Forest REU Site (McDevitt et al. 2016). Although we could predict difference in various learning gains based upon prior experiences, we had limited ability to explain the phenomena we observed. The design of our short self-reporting survey was an intentional compromise between sample size and survey depth, and such concessions contribute to a lack a general mechanistic understanding of outcomes of undergraduate research experiences (Beninson et al. 2011, Linn et al. 2015, Wilson et al. 2018). We hypothesize that the impediment for understanding why our, or other REU programs are successful is that our assessment tools were developed without an overarching theoretical framework. In this paper, we present a theoretical framework that we think would be useful for assessing and evaluating REU sites both singly and together.

## Using systems-based theoretical frameworks to study REU sites

Biologists have long recognized the complexity of biological systems and have developed techniques and models to study the interconnected components that make up these systems (Trewavas 2006). Systems-thinking similarly could be applied to understand and evaluate REU programs if relevant system components could be identified and adequately contextualized. In self-evaluative studies of single REU sites, researchers can provide some context about the design of the program and collect anecdotal or incidental evidence pertaining to individual participants, but the result still only is *post-hoc* speculation about the effectiveness of specific components of each program. This type of approach is common in discipline-based education research, which has a history of studies conducted in only one context (e.g., a single course, often that of the researcher) for which limited descriptions make generalization difficult (Singer et al. 2012). Although switching to a systems approach will not solve these problems completely, it can bring greater recognition to the interactions between system components that lead to site-specific outcomes and that may support broader generalizations.

Discipline-based education research is a relatively new area of research that continues to struggle to situate data within theoretical frameworks that include explicit statements of our theoretical assumptions with respect to learning. The extent to which studies are designed atheoretically varies by discipline, but biology-education research has been much more susceptible to this practice than the more established field of physics-education research (Singer et al. 2012). Lacking an explicit theoretical framework limits researchers’ ability to draw inferences and frames the research as observational or hypothesis-generating. Therefore, we recommend that biologists who are interested in education research begin by designing experiments in accordance with established educational theories (equivalent to scientific hypotheses).

### Cultural-historical activity theory (CHAT) as a broad theoretical framework

There are multiple theoretical frameworks that treat learning and learning environments as a system, including Bronfenbrenner’s bioecological theory of human development (Tudge et al. 2009), activity theory (Engeström et al. 1999), and cultural-historical activity theory (Roth and Lee 2007), which we discuss here. Although the scope of the research question should ultimately determine the selection of a theoretical framework (or competing frameworks), we think that cultural-historical activity theory (CHAT) is well-structured to inform the assessment of REU programs. CHAT is an expansion of activity theory (Engeström et al. 1999) that allows researchers to study the completion of goals either by individuals or collaborative groups while recognizing the cultural and historical influences acting on the system (Roth and Lee 2007).

CHAT provides a broad blueprint describing the components that influence the social construction of knowledge (Cole and Engeström 1993). This systems approach to research has proven useful for studying complex and heterogeneous educational phenomena and helps to derive meaning from seemingly contradictory information (Talbot et al. 2016). However for discipline-based education researchers, there are many barriers to using established theories, such as different approaches towards constructing knowledge, extensive educational jargon (Ravitch 2010), and disciplinary silos. To break down some of these barriers, we connect readers with some of the foundational literature from psychology, guiding questions to help interact with CHAT system components (Table S1), a simple rubric to help self-evaluate the strength of evidence for each CHAT system component (Table S2), and extensive examples for how to apply these resources during REU assessment (Table S3-S5).

Cultural-historical activity theory’s activity systems are best visualized through what are known as “activity triangles” (Roth and Lee 2007). CHAT requires the acknowledgement of seven distinct elements (“nodes”) that take part in an activity within a system of interest and the examination of connections (“edges”) between them (Roth and Lee 2007, Cole and Engeström 1993, Yamagata-Lynch 2010) (Figure 1). To help readers break down the educational jargon associated with CHAT, we illustrate its key pieces in describing a student writing a research proposal:

1. *Subject* – The individual or group of focus during the specified activity (e.g., the undergraduate student(s) writing the proposal);
2. *Object* – The goal or motive behind the specified activity (e.g., students should think critically about their project, connect with the primary literature, and establish feasible milestones for it);
3. *Rules* – The stated or unstated rules that govern how individuals act within the context of the specified activity (e.g., proposal guidelines, conventions of scientific writing, lab expectations or culture as established by research mentor);
4. *Community* – The social context in which the specified activity is conducted (e.g., including the student, research mentor, members of a lab, broader group of student participants);
5. *Division of labor* – How tasks are shared among the community to accomplish the specified activity (e.g., the student is responsible for most of the writing, the mentor provides some direction and feedback, and other lab members are available to answer questions);
6. *Mediating artifacts* – The tools used in creating or completing the *Object* (e.g., example project proposals, relevant journal articles, planning meetings, written feedback);
7. *Outcome* – The effect generated by subject working in concordance with other components of the activity system to accomplish the *Object* (e.g., formal evaluation of written proposal, performance review based on expectations outlined in proposal).

**Figure 1.**
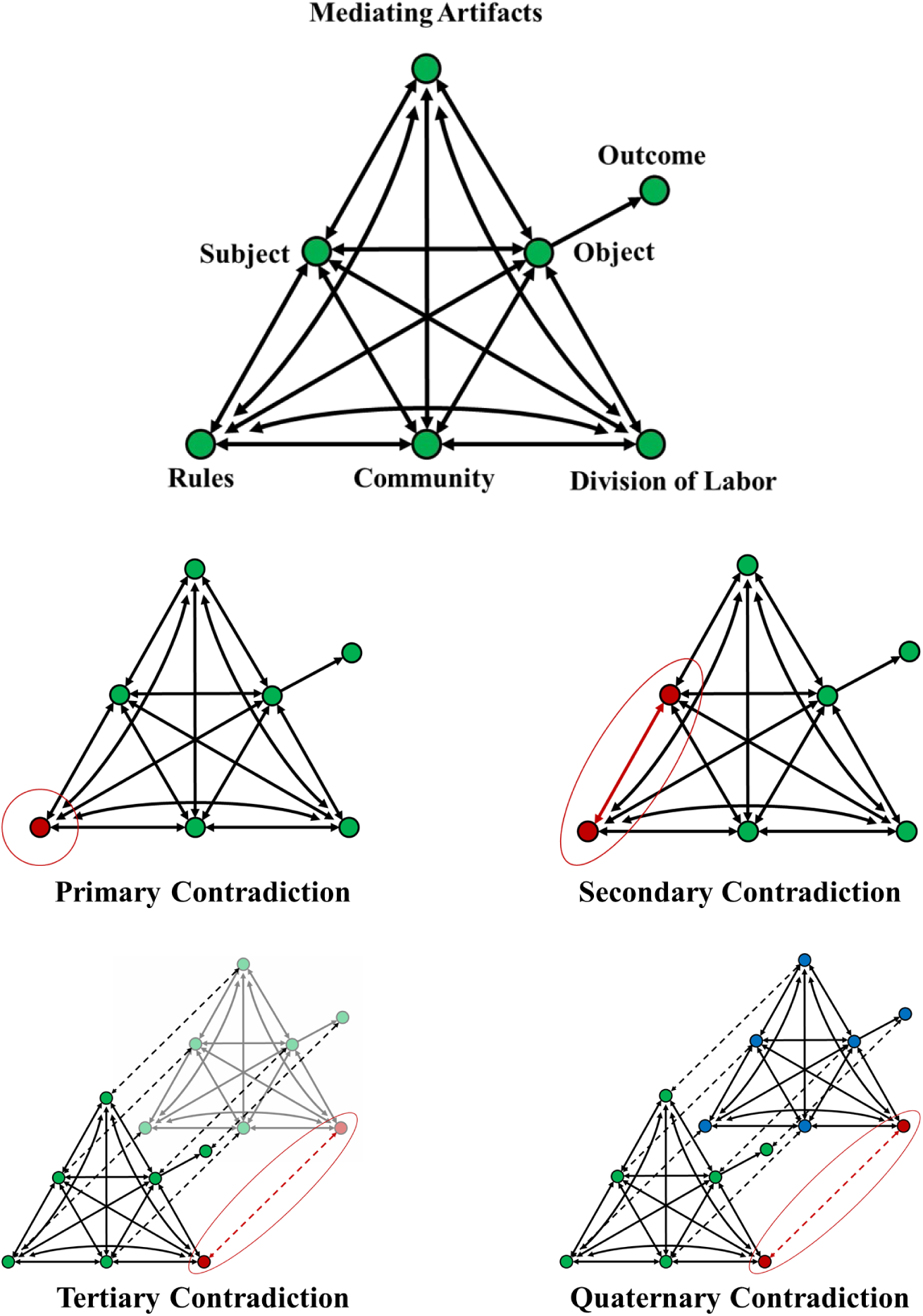
CHAT’s activity system components. The activity triangle highlights how components interact with others within the system (top), and the contradictions that can be examined through CHAT.

REU programs are complex social learning environments and CHAT provides the ability to make sense of contradictory information that arises within the system and through time (Cole and Engeström 1993). These contradictions, which often are difficult to rationalize using other frameworks, are classified into four types: *Primary contradictions* (1°) exist within an element *(e.g.*, contradictory *Rules*); *Secondary contradictions* (2°) exist within interactions between two elements (*e.g*., *Division of labor* is not aligned with *Mediating artifacts*); *Tertiary contradictions* (3°) are manifested during temporal transitions of an activity system (*e.g.*, mentors refining or modifying their approach “on the fly” while the student is writing their research proposal); and *Quaternary contradictions* (4°) that exist between similar activity systems of which the subject is a member (*e.g.*, REU experience compared to scientific coursework) (Engestrom 1987).

Primary contradictions within the CHAT framework often are a result of differing value judgements that underlie the system (Engestrom 1987). These contradictions are fundamental to the system and form the foundation of higher orders of contradictions (Engestrom 1987, Foot and Groleau 2011). After program values are established, components within an activity system should be aligned to aid the *Subject* in accomplishing the *Object*, measured by the *Outcome(s)*. Revisiting the research proposal example, this would mean that a student (*Subject*) is supported in a way that helps them write a successful research proposal (*Object*) that is measured by the expectations set by their research mentor or review panel (*Outcome*). However, it is common that two or more of these components are not aligned. Secondary contradictions help to illuminate this misalignment and may lead to subsequent changes within the activity system (Engestrom 1987).

For example, an undergraduate student (*Subject*) writing a research proposal (*Outcome*) may not possess the necessary background knowledge to read a highly technical literature review on their topic (*Mediating artifact*); the research mentor or other lab members (*Community*) may not have enough time to adequately support the student by answering questions and providing feedback (*Division of labor*); or expectations conveyed via a micromanagement approach (*Rules*) conflict with the ability for the student to meaningfully connect with the literature or think independently about their project (*Object*). These conflicts between system components may result in specific obstacles that are manifestations of fundamental tensions *(Primary contradictions)* within the activity system (Foot and Groleau 2011). Because conflicts and contradictions may arise from fundamental components of the system, it is often better to address their source(s) rather than their symptoms. To resolve *Secondary contradictions* by addressing underlying *Primary contradictions*, some type of change must occur in the activity system. For example, before trying to develop new *Mediating artifacts* to help a student read a highly technical literature review (*Secondary contradiction*), it would be prudent to first evaluate if there already are M*ediating artifacts* in place that send conflicting messages (*Primary contradiction*), which once addressed, might resolve the *secondary contradiction*.

*Tertiary contradictions* are differences in the system that occur at temporal transitions (Engestrom 1987), and program directors may be interested in examining them as they change various instructional activities or procedures. For example, an REU program may implement a new proposal writing workshop (*Mediating artifact*) that is intended to help students (*Subject*) connect their proposals to the available scientific literature (*Outcome*) and simultaneously shift some of the duties from the research mentor to the workshop facilitator and the student’s peers (*Division of labor).* As new procedures are implemented, a transition to more “advanced” practices may not be immediate (Engestrom 1987, Foot and Groleau 2011). Examining barriers to change may reveal additional information about *Primary contradiction*s and potentially lead to smoother tertiary transitions.

Alternatively, the cause of these underlying contradictions may not reside solely within the activity system itself, but rather may rooted in cultural expectations from adjacent activity systems (*Quaternary contradictions*). Students (*Subjects*) brings their past experiences with them to the activity system, and it is likely that members of the *Community* may not have the same shared experiences. For example, the *Rules* established in adjacent activity systems may carry over for an individual and impact how they interact with system components such as *Mediating artifacts* or the *Community.* For example, if a student (*Subject*) has prior experience writing a research proposal (*Object*) in another context (e.g., in a different lab, discipline, or institution), their perceptions of this current experience in writing may be influenced by *Rules, Mediating artifacts*, or *Division of labor* from their other experience (*Adjacent activity system*). In this case, the success in writing their REU research proposal (*Outcome*) is driven by the recognition of these *Quaternary contradictions* and relevant interventions, such as the adjustment of *Rules*, addition of *Mediating artifacts*, or changes to the *Division of labor* that can lead to more productive writing process by the student (*Subject*).

## Applying the CHAT framework to REU research

Harvard University established its Summer Research Program in Ecology at the Harvard Forest (HF-SRPE) in 1985; NSF has provided continuous core support for the HF-SRPE through REU Site awards since 1989. Through three decades, we and our predecessor PI and co-PIs have participated in the development and implementation of NSF’s vision for REU sites. Simultaneously, we have enhanced student experiences and improved short- and long-term effectiveness of our program by regularly measuring and reflecting on its success and failures and integrating our assessment data with REU-wide evaluations (McDevitt et al. 2016). Based on our experiences, we advocate for integration of theoretical frameworks such as CHAT in assessment and evaluation to increase the efficiency by which REU sites use individual and cross-site assessment data to improve their own programs. Below, we provide three examples by which CHAT can be applied to contextualize HF-SRPE and improve our assessment protocols. We provide rubrics aligned with CHAT (Tables S1–S2) to guide assessment-design decisions and HF-SRPE responses for three activity systems of interest (Table S3–S5). In doing so, we aim to increase dialogue and the sharing of administrative procedures which we hope will strengthen educational research and ultimately the experience of participants.

### Recruitment and hiring practices

The first stage of HF-SRPE and any REU program is to recruit and hire student participants (Table S1). In the language of CHAT, the *Activity triangle* (Figure 1) represents the recruitment process. We use characteristics about the individual (*Subject*), the priorities of the position (*Object*), the expectations of the hiring process (*Rules*), and practices implemented in recruiting or selection (*Mediating artifacts*) to recruit and hire a diverse set of students with various degrees of prior experience participating in mentored research (*Outcome*).

At these earliest stages of the program, a *Primary contradiction* exists in the activity. We try to strike a balance between selecting students who appear to be best qualified (i.e., most experienced) to conduct research and those who have the most to gain out of the experience. This contradiction arises in part from cultural biases of academic research where success is measured through productivity (theses, posters, peer-reviewed papers); the “best” students are those with proven “track records” of productivity. However, as mentors and educators, we also want to work with students who are willing to push beyond their comfort zone and maximize the impact of a research experience. At HF-SRPE, this *Primary contradiction* is further complicated by the different stakeholders involved in the hiring process. Individual research mentors advocate for their projects; funders push for students from certain institutions, demographics, academic majors, or skillsets; and program directors seek a lasting and cohesive identity for the program. Characterizing these various components and assessing if recruitment and hiring goals are being met is difficult, especially when hundreds-to-thousands of applications are reviewed in a scant few weeks.

Demographic data and quantifiable metrics such as gender, ethnicity, grade-point average (GPA), class rank, or type of institution may be the easiest variables for a single site to gather about its applicants, to use in decisions about who to interview or hire, and to track through time or compare among multiple REU sites. However, for REU programs to evaluate the effectiveness of practices such as recruitment, application requirements, and selection criteria, the data collected need to be aligned with the broader context and goals of the activity (mentored research at an REU site). This is where a meta-theoretical framework like CHAT can be used to identify and prioritize useful data to be collected (Figure 2). Although it would be best to collect lines of evidence supporting each CHAT component, priorities for HF-SRPE (Table S3) at a minimum would be to collect more informative characteristics about individual applicants (*Subjects*), the priorities of the mentors filling each position (*Object*), the expectations of the site PIs in recruiting a diverse group of participants (*Rules*), recruiting and selection (*Mediating artifacts*), and the final hiring decision (*Outcome*). Rather than serving as a predictive theory about the hiring process (which could alternatively be integrated into an assessment program), CHAT provides focus and clarity for understanding this complex system.

**Figure 2.**
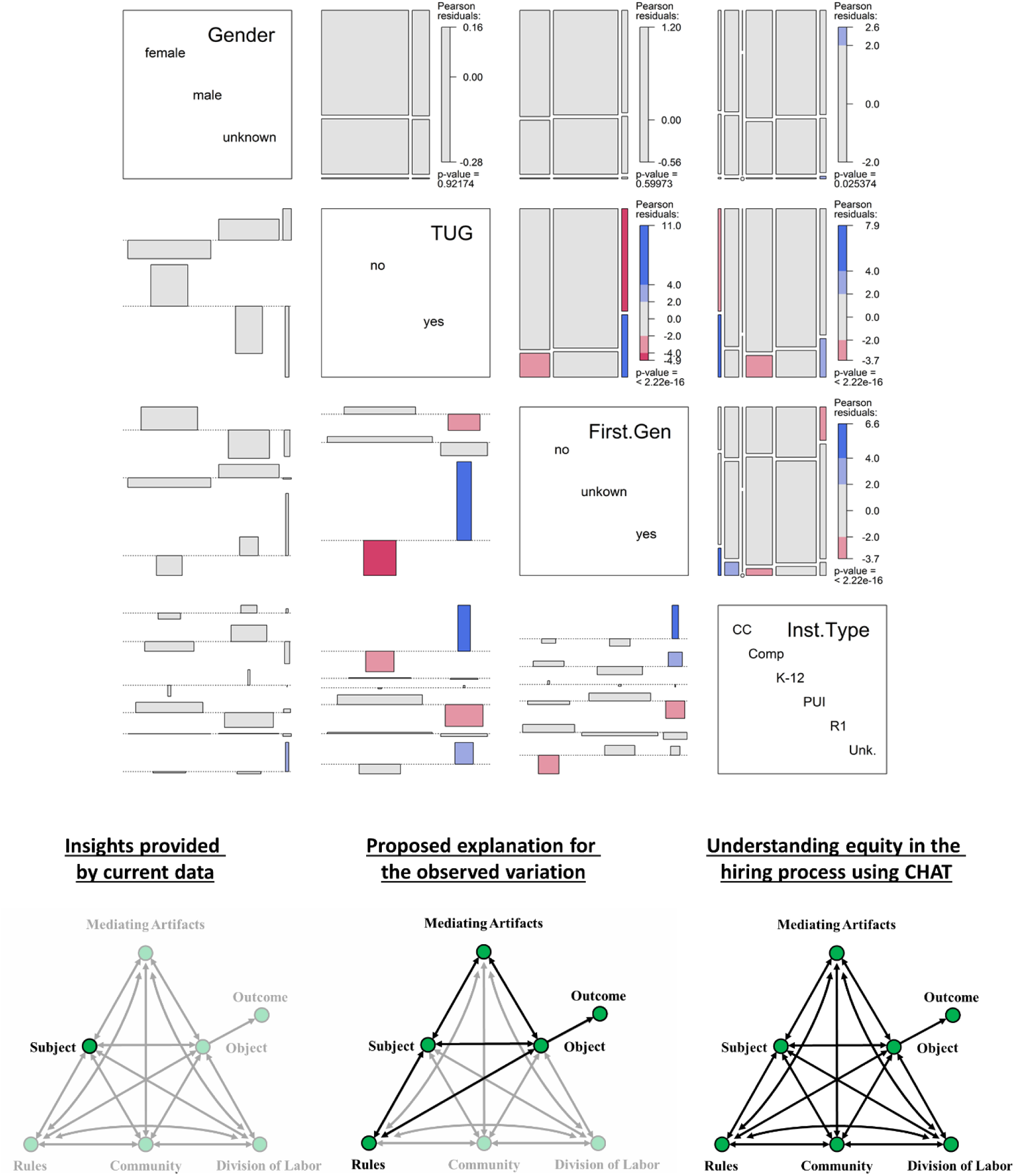
Data that are commonly collected during the recruitment and hiring process (top) and the CHAT activity triangles illustrating how components could be assessed with current frameworks (bottom left) or could be assessed within a full CHAT framework (bottom center and bottom right). The top panels show mosaic plots illustrating relative proportions of students in different groups of quantitative interest to students, program directors, programs, and funders. Significant positive (blue) and negative (red) pairwise correlations are indicated. TUG: Student from groups traditionally under-represented in science; First.Gen: Students who are the first in their family to attend college or university; Inst.Type: type of institution, including community college (CC), Comprehensive university (Comp), K-12 (kindergarten through high school), PUI (primarily undergraduate institution), R1 (research-1 university), and Unk (unknown or not applicable).

### Understanding variation in learning gains

The scientific theme for the most recent five-year (2015-2019) REU Site award for HF-SRPE has been the collection, visualization, analysis, and communication of ecological “Big Data.” Like other REU sites in biology, we have used URSSA to provide self-assessment of gains in learning through questions about broad items related to thinking and working like a scientist (Hunter et al. 2009). URSSA includes questions that address students’ attitudes, feelings, and motivation related to analyzing data for patterns, problem solving, and identifying limitations. Superficially, these may seem like they can assess the learning gains of interest, but the developers of URSSA defined its scope only as a broad indicator of progress (Weston and Laursen 2015). The questions are not aligned with our specific program goals (i.e., poor criterion validity) and are unable to provide meaningful measurements for any of our “Big Data” learning outcomes. Additionally, the limited student or programmatic context provided by URSSA, such as demographics (Figure 3), rarely accounts for much of the variation in URSSA’s measured gains. Such limitations have constrained our ability to make informed decisions for program improvement or providing evidence that HF-SRPE is helping students achieve these goals.

**Figure 3.**
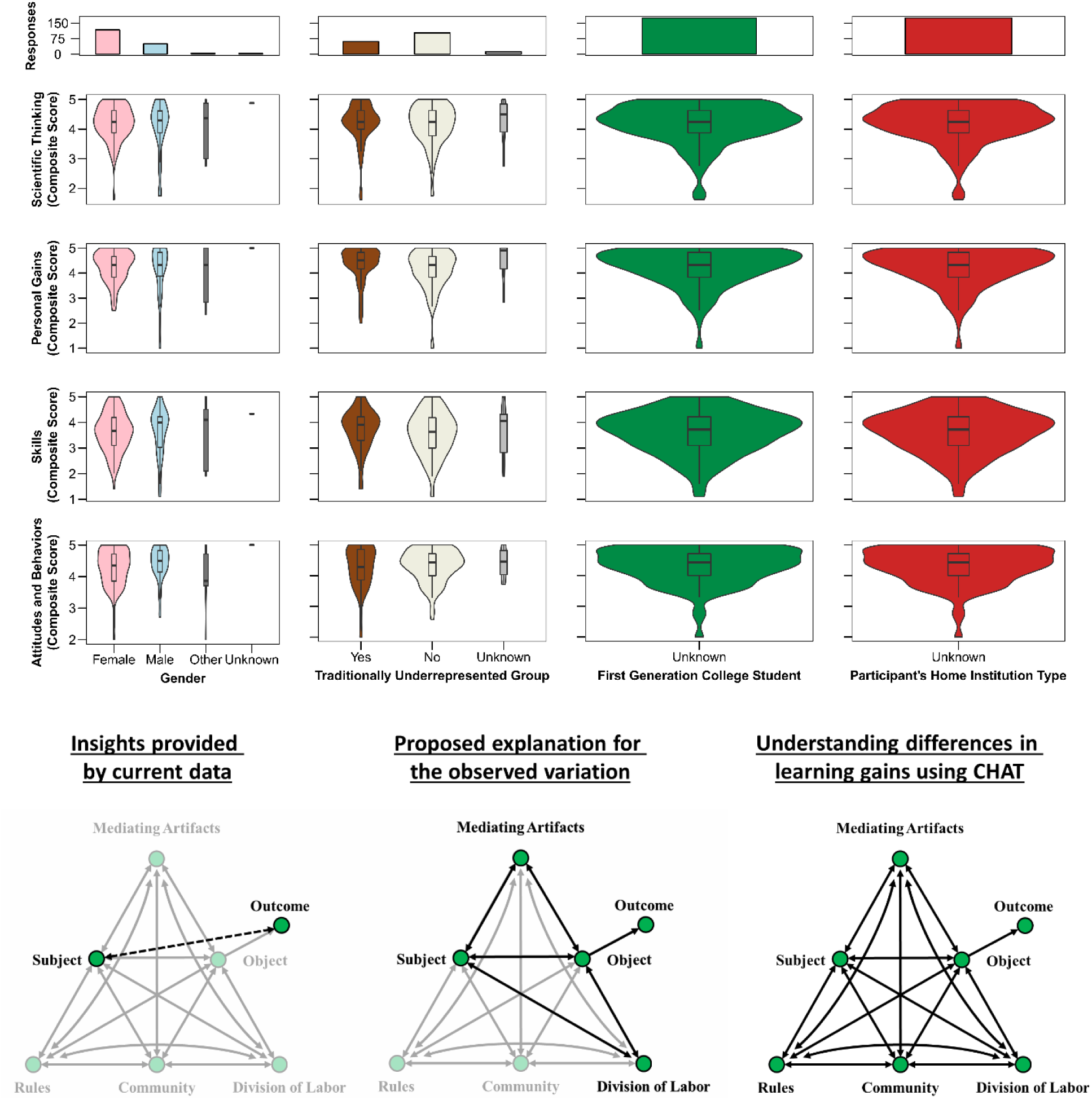
Data commonly collected when assessing learning gains (top) and CHAT activity triangles illustrating how components could be assessed with current frameworks (bottom left) or within a full CHAT framework (bottom center and bottom right). The top panels show changes in scientific thinking, personal gains in overall confidence in doing research, research skills, and attitudes and behaviors about doing research. Values range from 1 (low) to 5 (high) for all variables. The total number of participants in the different groups are shown in the top row; in the other panels, violin plots show the distribution of the data with inset box plots illustrating median, quartile, and upper and lower deciles of the data. More detailed analysis of these data, collected from pre/post surveys given annually to HF-SRPE students, are presented in Weston and Laursen (2015).

Alternatively, we have used our knowledge of HF-SRPE to describe components and contradictions within a CHAT activity system (Table S1) to identify more useful data to address our learning objectives. Although it would be ideal to align and characterize all seven components of the activity system with respect to learning gains, we also need to set priorities for assessment. To do so, we examined our response to Table S1, evaluated the quality of current (or planned) evidence, and made judgments about data to collect based on our knowledge of the literature and through running HF-SRPE (Table S4). This exercise led us to prioritize the following aspects for explaining the variation in student learning gains (Figure 3): characterizing the skills and knowledge a student brings with them to the research experience (*Subject*); the resources available to the student during their research experience (*Mediating artifacts* such as R workshops and project proposals); the level of support they received (*Division of labor*); and what success means given a student’s prior research experience (*Object*). This richer characterization of the REU learning environment would then be used to help explain differences in learning gains (*Outcome*).

We would ideally assess learning gains (*Outcome*) using a concept inventory (i.e. a validated educational instrument that allow educators to evaluate student ideas and beliefs about a topic) related to our “Big Data” learning objectives. Although concept inventories often are praised for their extensive reliability and validity testing, they usually are designed to assess specific concepts most frequently obtained through classroom coursework. Rarely are concept inventories aligned with the objectives of experiential learning that occurs in REU programs. To assess our “Big Data” learning objectives in an REU context would require the development and refinement of a new concept inventory (developing what is colloquially known as a “million-dollar instrument”). In the meantime, learning objectives can be assessed through student self-evaluations (recognizing that students are likely to have difficulty evaluating a topic they are not yet proficient), mentors’ evaluations of student proficiencies (that will also depend on the mentors’ proficiency and their familiarity with students’ progress), or a more detailed analysis of research products (that may not represent the breadth of what a student learned).

### Assessing the impact of REU programs

Qualitative feedback from previous REU participants have suggested that mentored independent research is a formative experience for their career development. Systematic, post-program tracking of REU participants is uncommon despite it being a legal requirement since 2010 (P.L. 111-478 §514). We annually survey past participants of HF-SRPE; the resulting data provide some support for long-term persistence and high rates of employment by HF-SRPE alumnae/i in STEM fields (Figure 4). However, we have been unable to account accurately for the distribution of non-responses or determine specific effects of HF-SRPE on individual decisions to pursue STEM careers. Like many of our colleagues who work with REU students, we believe that mentored research experiences launch them into STEM careers, but we cannot predict where they would be without this experience. Ethical and logistical constraints prevent researchers from forming true control groups for REU participants; we can use only quasi-experimental designs. Expansions of data collection efforts before and during REU programs are needed to meaningfully characterize students and their experiences.

**Figure 4.**
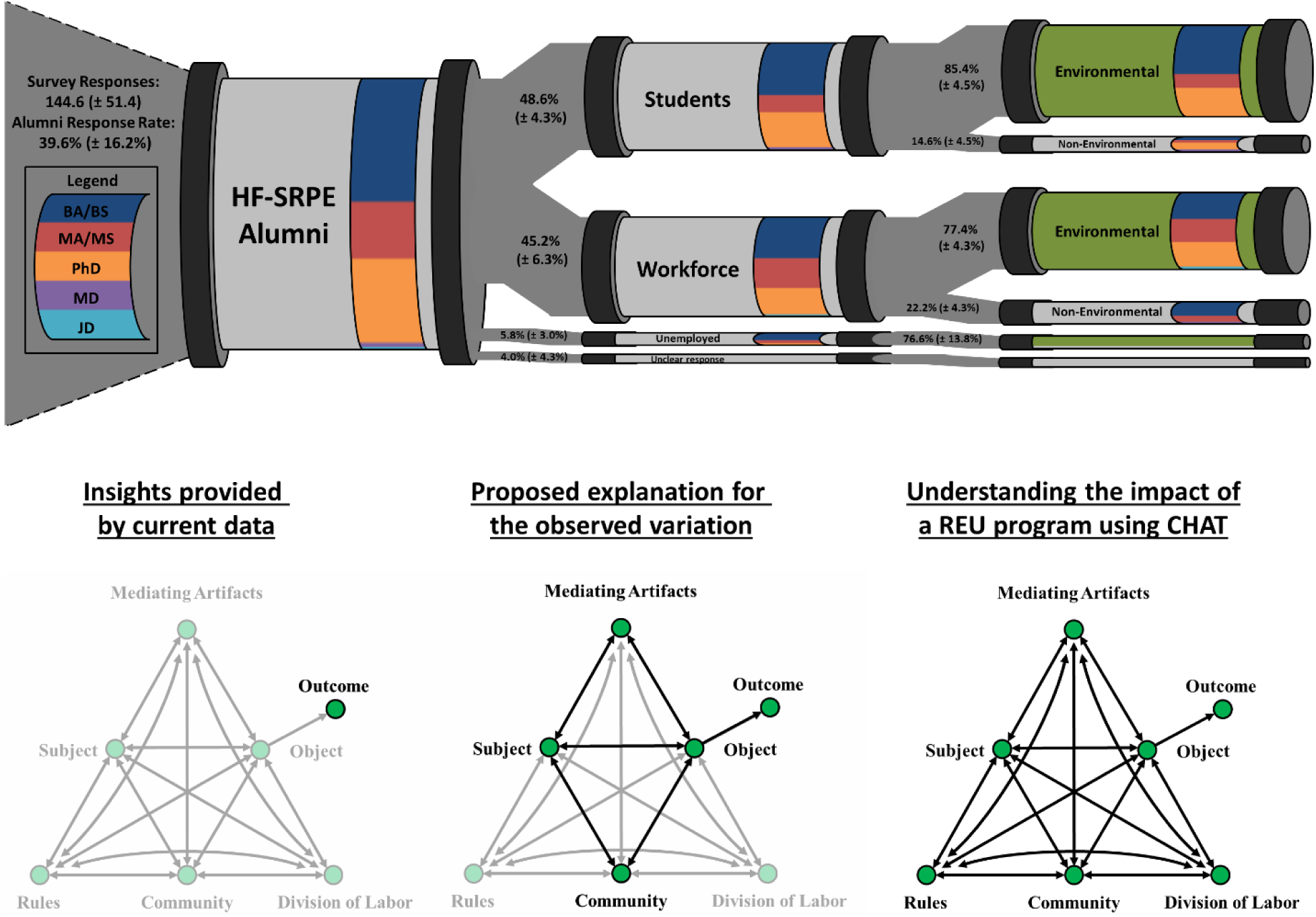
HF-SRPE career outcomes (top “pipeline”). Annual alumni surveys were sent to alumnae/i (cohorts from 2001 onward) between 2012 and 2016. Averages of yearly snapshots reveal that alumnae/i have pursued or received environmental- or ecology-related graduate degrees and continue to use these disciplines during their careers. Further information is required to determine the impact of HF-SRPE on these outcomes. The CHAT activity triangles (bottom) illustrate how components could be assessed with current frameworks (bottom left) or within a full CHAT framework (bottom center, bottom right).

The systems approach of CHAT can be used to identify important aspects of the complex learning environment and be used to generate more specific hypotheses about how REU programs impact the long-term persistence of participant in STEM disciplines or careers. Using the CHAT worksheets (Tables S1–S2), we would prioritize collection of the following data about HF-SRPE: the skills and knowledge a student brings with them to the research program (*Subject*); the professional development opportunities available to them during their research experience (*Mediating artifacts*); the interactions students have with other members of the research community (*Community*); and the goals of the research experience (*Object*) (Table S5).

We admittedly know little about why our students do or do not persist in STEM disciplines or careers after HF-SRPE. There has been evidence to suggest students participating in structured programs obtain advanced degrees and generate research products at a higher rate compared to pair-matched students who applied but were not selected (Wilson et al. 2018). If we were to replicate this experiment with the addtion of CHAT, we might consider the inclusion of additional working hypotheses (Chamberlin 1890) such as the *Rules* governing why students selected to participate (which may be independent of basic demographic descriptors like gender, ethnicity, home institution, or GPA), the specific *Mediating artifacts* used to help students achieve their *Object* (recognizing that there are likely multiple equivalent paths to long-term success), and acknowledging how students (*Subjects*) and other members of their *Community* may view success (*Outcome*).

## Conclusions

Systems-based theoretical frameworks such as CHAT can help to identify and coordinate the type of information that should be collected so that experiential research and education programs can accurately assess how well they are achieving their goals and knowledge can be transferred more easily between programs. As with most scientific inquiry, the research questions ultimately should drive the types of data that are collected. As REU sites are developing assessment protocols, it would be beneficial to collect data within the framework of multiple competing hypotheses (Chamberlin 1890). We have provided guiding questions to do so using CHAT (Table S1) and would recommend similar exercises if other theoretical frameworks were to be applied in assessing undergraduate research programs.

Selecting the proper framework requires both a clear understanding of programmatic goals and a familiarity with the theory and literature in education research. The latter is particularly challenging for REU program directors who have not studied education research methods. We encourage those who may feel daunted by learning another discipline to seek collaborators in education or qualitative research while expanding one’s own fluency. However, once a proper framework has been identified, it should be easier to transfer meaningful experiences and best practices across the larger REU community and others working in similar educational contexts. The short-term nature and finite number of REU sites highlights the need to evaluate what works—and what doesn’t—and disseminate this information to the broader scientific community, lest we keep reinventing pedagogical wheels. As evaluative research continues to develop within the REU community, we see systems-based theoretical frameworks as useful guidelines for programs to follow when assessing REU programs.

## Acknowledgments

This work was supported by NSF grants DBI 04-52254, 10-03938, and 14-59519; and NASA GCCE award NNX10AT52A, all to AME. Data reported in this paper are available from the Harvard Forest Data Archive (http://harvardforest.fas.harvard.edu/data-archive/), datasets HF279, 280, and 281.

---

**Figure S1.**
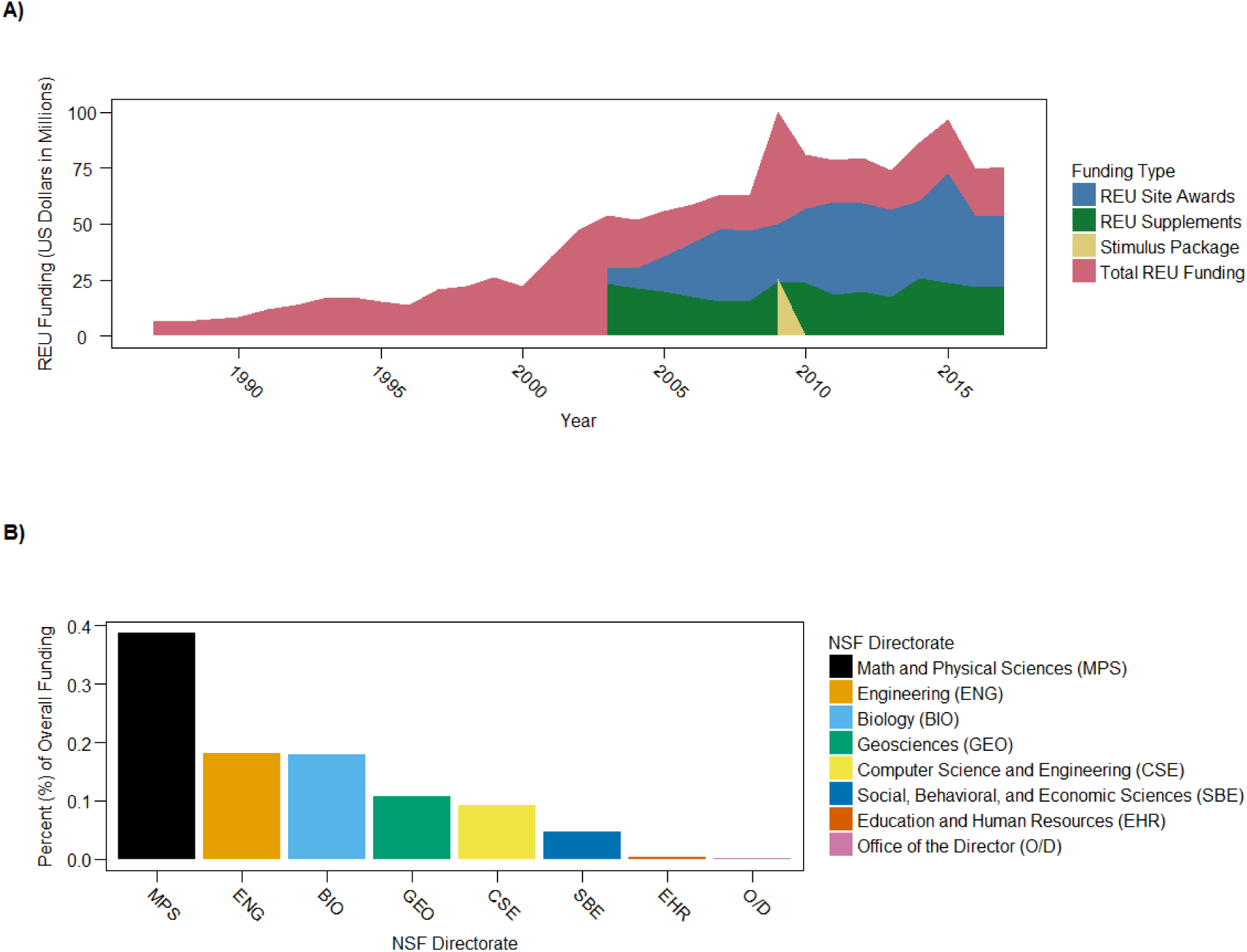
Funding for Research Experiences for Undergraduates (REU) programs. Support for REU programs based on A) yearly congressional allocations and B) NSF directorate support for REU Sites. Funding data (2002-2017) was compiled based on yearly NSF congressional budget requests. Archives of REU awards (nsf.gov/awardsearch/) provided estimates for remaining years and directorate contributions.

**Table S1.** Recommended questions for characterizing CHAT system components.

| System Component | Identifying Components | Identifying Contradictions |
| --- | --- | --- |
| Subject | **S.0** – As the evaluator, how would you characterize the subject(s)?<br>**S.0** - How might the subjects characterize themselves? | **S.1** - Is there a consensus about the characterization of the subject? What type of information might be relevant to the activity system, and how might you go about collecting this description?<br>**S.2** - See Below. Secondary contradictions will be examined once other components of the activity system have been examined.<br>**S.3** - If you are studying this activity system over time, how has the characterization of your subject changed over time?<br>**S.4** - What are some similar activity systems the subject may be involved in? |
| Object | **Ob.0** - What is the activity/practice you want to evaluate?<br>**Ob.0** - How would the subject characterize the object?<br>**Ob.0** - Are there other members of the community that would characterize the object differently? | **Ob.1** - Is there a clear consensus about the goals of the activity/practice under evaluation? If not, where might you expect to find additional contradictions?<br>**Ob.2** - How does the subject perceive the object? (e.g. Do they have the same priorities?)<br>**Ob.3** - If you are studying this activity system over time, how has the object changed over time?<br>**Ob.4** - What are additional activity systems might have similar objects/goals? |
| Outcome | **Out.0** - What values or perspectives are important to recognize when assessing the object? (e.g. student “success” may mean many things to different people)<br>**Out.0** - How do you plan on assessing the object? | **Out.1** - Why is this outcome measurement appropriate for the object? What are the documented validity and reliability arguments for this object?<br>**Out.2** - What evidence do you have that the outcome measures are aligned with your object?<br>**Out.3** - If you are studying this activity system over time, how have your outcome measures changed? (e.g. refinement or development of instruments/surveys)<br>**Out.4** - If you are using an instrument that was developed to under a different context/population, how might this impact the validity and reliability arguments? |
| Community | **C.0** - Who interacts with the subject to accomplish the object? | **C.1** - Does the subject acknowledge or recognize the rest of the members of the community?<br>**C.2** - Do both the subject and community know who is involved in the activity system?<br>**C.3** - If you are studying this activity system over time, how has the community changed?<br>**C.4** - How does this community compare to similar activity systems? What is the impact these differences have on the outcome? |
| Rules | **R.0** - What rules, expectations, or cultural norms are present that potentially impact how the subject interacts with the activity system? | **R.1** - Are any rules or norms at odds with one another? (i.e. can they send conflicting messages)<br>**R.2** - Are any rules in direct conflict with the object?<br>**R.2** - Do subjects or community members all value or agree with the rules of the system?<br>**R.3** - If you are studying this activity system over time, how have rules changed over time?<br>**R.4** - How do the rules of this activity system differ from similar activity systems? What impact do these differences have on the outcome? |
| Mediating Artifacts | **MA.0** - What tools or practices are used help the subject achieve/accomplish the object? | **MA.1** - What evidence can be provided there that the various mediating artifacts are aligned?<br>**MA.2** - How does the use of mediating artifacts align with the object?<br>**MA.2** - How does the subject perceive the utility of individual mediating artifacts?<br>**MA.2** - How does the community perceive the utility of individual mediating artifacts?<br>**MA.2** - Are any rules in conflict with mediating artifacts (or with how the community divides the labor with these artifacts)?<br>**MA.3** - How did the addition, removal, or refinement of mediating artifacts impact the activity system?<br>**MA.4** - How do these mediating artifacts compare to similar activity systems? What is the impact these differences have on the outcome? |
| Division of Labor | **DL.0** - What are the expectations of the community to be involved in helping achieve the object? | **DL.1** - What evidence can be provided to demonstrate that community members are achieving these expectations?<br>**DL.2** - Is the division of labor appropriate to meaningfully support individual mediating artifacts?<br>**DL.2** - Is the overall division of labor in the activity system appropriate to meaningfully support the object?<br>**DL.3** - If you are studying this activity system over time, how has the division of labor changed over time?<br>**DL.4** - How does the division of labor differ from similar activity systems? What is the impact these differences have on the outcome? |

**Table S2.** Recommendations for rating the quality of evidence in CHAT systems.

| <u>Measurement</u> |  | <u>Assessment Design</u> |  |
| --- | --- | --- | --- |
| Qualitative data | Quantitative data | Alignment | Sampling |
| **QL.0**<br>Data have not been collected. | **QT.0**<br>Data have not been collected. | **A.0**<br>Data and question are clearly unaligned, or no data has been collected | **S.0**<br>Data have not been collected. Information is solely based on researchers' perceptions. |
| **QL.1**<br>Data have been collected. However, construct validity has not been formally evaluated or analyses were done incorrectly. | **QT.1**<br>Data have been collected. However, construct validity has not been formally evaluated or analyses were done incorrectly. | **A.1**<br>Data have face validity. Criterion validity has not been formally evaluated or analyses were done incorrectly. | **S.1**<br>Data were collected through convenience sampling. Helps generate hypotheses but inferences cannot be made beyond sample. |
| **QL.2**<br>Formal evaluations of construct validity have been conducted. However, evidence for both internal or external validity is limited, or studies were not conducted in your context of interest | **QT.2**<br>Formal evaluations of construct validity have been conducted. However, evidence for both internal or external validity is limited, or studies were not conducted in your context of interest | **A.2**<br>Formal evaluations have been conducted and provides some evidence for criterion validity. | **S.2**<br>Data were collected intentionally. However, sampling may have occurred at an incorrect scale, there is sampling bias, or participants choose not to participate. |
| **QL.3**<br>Formal evaluations of construct validity have been conducted. There is strong evidence for both internal or external validity in your context of interest | **QT.3**<br>Formal evaluations of construct validity have been conducted. There is strong evidence for both internal or external validity in your context of interest | **A.3**<br>Formal evaluations have been conducted and provides strong evidence for criterion validity. | **S.3**<br>Data were collected using a random sample of the population (or sample frame) of interest or data exists for the entire population. |

**Table S3.** Example responses to the CHAT questionnaire (Table S1) for assessing participant selection at HF-SRPE.

| Question<br>(Table S1) | Response | Quality of Evidence<br>(Table S.2) | Relative priority to address<br>(Low, Med, High) |
| --- | --- | --- | --- |
| S.0 | HF-SRPE is a paid summer research program for undergraduate participants located at the Harvard Forest in Petersham, Massachusetts. Over the past 30 years HF-SRPE has been primarily supported through the NSF REU program, however additional participants within the program have been supported through various grants and programs. | QL.1<br>QT.0<br>A.1<br>S.1 | High |
| S.0 | Same as above. This is an example of self-evaluation. | QL.1<br>QT.0<br>A.1<br>S.0 | Low |
| S.1 | Yes. However, since this is a self-evaluation it will be important to record the relevant context so that others can adequately understand the nuances of the activity system. | QL.1<br>QT.1<br>A.1<br>S.1 | High |
| S.3 | There have been subtle changes in program structure and funding over the past 30 years. We have an archive of general descriptions of the program through grant documents and advertising material, however a formal document analysis has not been conducted to address this question. | QL.1<br>QT.0<br>A.1<br>S.1 | Med |
| S.4 | Other REU programs and field ecology research experiences; Harvard College admissions. The applicants and mentors may perceive aspects of this activity system differently based on their experiences or knowledge of similar *Activity Systems*. | QL.0<br>QT.0<br>A.0<br>S.0 | Med |
| Ob.0 | Select participants for the Harvard Forest Summer Program in Ecology | NA | NA |
| Ob.0 | Equitable hiring process. Support interests such as providing research opportunities to undergraduate students who can benefit the most from the program, supporting research programs of PIs, the mission of the Harvard Forest | QL.1<br>QT.0<br>A.1<br>S.1 | High |
| Ob.0 | Mentors may have similar perceptions of the selection process. However, they could possibly prioritize aspects such as speed or ease of hiring process, demonstrated skills of applicant (as opposed to potential. This has not been examined directly and would warrant further investigation. | QL.1<br>QT.0<br>A.1<br>S.1 | High |
| Ob.1 | Overall there seems to be alignment, however with many stakeholders there may be conflicting values and expectations (rules) | QL.1<br>QT.0<br>A.1<br>S.1 | High |
| Ob.2 | This is a self-evaluation of HF-SRPE and are doing our best at introspection. Individuals within the *community* may have different priorities (see below). | QL.0<br>QT.0<br>A.0<br>S.0 | Low |
| Ob.3 | The need to select participants for the program has remained the same (although values, rules, mediation artifacts, etc. have changed; see below) | QL.0<br>QT.0<br>A.0<br>S.0 | Low |
| Ob.4 | Undergraduate admissions, graduate admissions, other REU programs:<br>Members of the *Community* may perceive aspects of this activity system differently based on their experiences/knowledge of similar *Activity Systems* | QL.0<br>QT.0<br>A.0<br>S.0 | Low-Med |
| Out.0 | HF-SRPE wishes to support equitable hiring practices, promote diversity in the field ecology, and select participants who can benefit the most from the program. | QL.1<br>QT.0<br>A.1<br>S.1 | High |
| Out.0 | Primarily only demographic information (gender, ethnicity, institution type, first generation status) | QL.1<br>QT.1<br>A.1<br>S.3 | High |
| Out.1 | We haven't identified any contradictions between the measurement and our values associated with the hiring process. | QL.0<br>QT.0<br>A.0<br>S.0 | Med |
| Out.2 | The demographic information is broadly aligned with most of the goals of the program. However, with regards to inclusive hiring practices, these data only provide a little bit of insight into an applicant's personal identity. | QL.0<br>QT.0<br>A.0<br>S.0 | Med |
| Out.3 | Due to the simplicity of these items, most of these measurements have not changed | QL.0<br>QT.0<br>A.0<br>S.0 | Low |
| Out.4 | The data we collect is very similar to that of adjacent *activity systems*. While we imagine | QL.0<br>QT.0<br>A.0<br>S.0 | Low |
| C.0 | Undergraduate applicants, Principle Investigators, HF-SRPE staff, HR | QL.4<br>QT.1<br>A.1<br>S.4 | High |
| C.1 | Mostly, however HF-SRPE has a difficult time identifying participants who do not apply to the program (but might otherwise turn out to be competitive applicants). Without knowing who those individuals are, or their barriers to application, HF-SRPE is limited in its ability to determine if it is providing an equitable application process. | QL.?<br>QT.0<br>A.?<br>S.0 | Med-High |
| C.2 | This question is targeted to *Subjects* who are individuals and is an awkward to ask of a program in self-evaluation. | NA | NA |
| C.3 | It is often different each year, with overlap in admin staff and mentors. Sometimes there are repeat applicants and references. | QL.0<br>QT.1<br>A.1<br>S.3 | Med |
| C.4 | We have anecdotal evidence that our *Community* is similar in structure to that of other REU programs. However, we do expect that the size of our applicant pool (~800 undergraduate annually), PIs, and program staff are greater than most REU sites. | QL.1<br>QT.1<br>A.1<br>S.1 | Med-High |
| R.0 | Rules and expectations of this hiring process are akin to most other employment opportunities. Although mentors can set their own expectations for hiring criteria, HF-SRPE program staff try to unify program expectations (e.g. projects involving intellectual participation of the participant, sharing equitable hiring | QL.0<br>QT.0<br>A.0 | High |
|  | resources, communicating expectations of funding agencies). HF-SRPE also provides logistical advice for applicants to clarify program expectations for both the application process and the summer experience. | S.0 |  |
| R.1 | We have tried to be consistent in advertising material, hiring expectations, and values. However, we have not conducted formal evaluations to see if the messages we send are aligned with our values (e.g. explicitly review all advertising material and requirements through a multicultural lens) | QL.0<br>QT.0<br>A.0<br>S.0 | High |
| R.2 | There do not appear to be any rules or admission requirements that would limit us from hiring participants for all available positions. | QL.0<br>QT.0<br>A.0<br>S.0 | Low |
| R.2 | *Community* members within the activity system may have conflicts with the rules, expectations, and cultural norms of the hiring process. Mentors, who ultimately do the hiring, may have different priorities or values compared to what is stated (or assumed) by the program. Applicants may also have different priorities or values, especially qualified students we are failing to reach or recruit. | QL.0<br>QT.0<br>A.0<br>S.0 | Med-High |
| R.3 | Rules, deadlines, and expectations are routinely changing as we reflect more on the hiring process. For example, over the past 15-20 years we have increased our commitment to recruiting a diverse applicant pool. These values have helped shape improvements to mediating artifacts such as recruitment methods and the hiring process. | QL.0<br>QT.0<br>A.0<br>S.0 | Med-High |
| R.4 | We believe that HF-SRPE's rules and expectations are comparable to other REU programs or internship programs. However, it is possible that there are cultural norms establish by other programs that are in contrast with HF-SRPE and may be a barrier to recruiting certain qualified participants. | QL.0<br>QT.0<br>A.0<br>S.0 | Med |
| MA.0 | Application materials (position advertisements, program website)<br>Submission tool (front-end used by applicants and references to submit application materials)<br>Submission tool (back-end used by mentors and HF-SRPE staff to review applicants)<br>Hiring documents (project descriptions, interview protocols, expectations) | QL.1<br>QT.0<br>A.1<br>S.2 | High |
| MA.1 | These *Mediating artifacts* have undergone an iterative design process over many years, however a formal analysis has not been conducted to see if the mediating artifacts are aligned with each other. | QL.1<br>QT.0<br>A.1<br>S.1 | High |
| MA.2 | These *Mediating artifacts* have undergone an iterative design process over many years, however a formal analysis has not been conducted to see if the mediating artifacts are aligned with the *Object*. | QL.0<br>QT.0<br>A.0 | Low-Med |
|  |  | S.0 |  |
| MA.2 | This question is targeted to *subjects* that are individuals and is an awkward to ask of a program in self-evaluation. | NA | NA |
| MA.2 | In general, we believe that members of the *community* are finding the *Mediating artifacts* useful. Over the years, we have received informal feedback which has guided the design of our submission tools and hiring procedures. However, we are still uncertain how our recruiting process/materials might be limiting otherwise qualified applicants from applying. | QL.1<br>QT.0<br>A.1<br>S.1 | Med |
| MA.2 | We have not received feedback indicating that our *Rules* (or *Division of labor*) conflict with any of the *mediating artifacts*. | QL.0<br>QT.0<br>A.0<br>S.0 | Low-Med |
| MA.3 | We believe that the introduction of certain *Mediating artifacts* (e.g. online submission tool) have been beneficial to improving *outcomes* such as diversity in hiring. | QL.0<br>QT.0<br>A.0<br>S.0 | Med-High |
| MA.4 | Due to the high volume of applications (>500 per summer) and the digital infrastructure already in place at the Harvard Forest, HF-SRPE has invested a fair amount of effort into our online submission tool. We imagine that smaller (or newer) programs do not have the need for such infrastructure. While we feel that these mediating artifacts are important for HF-SRPE to achieve our goals, we anticipate that there are many successful alternatives | QL.1<br>QT.0<br>A.1<br>S.1 | Med-High |
| DL.0 | Application materials: HF-SRPE staff create and disperse most of these materials; mentors may also disperse materials through their own networks; Some applicants may need to seek out these materials, while others may be introduced to them through their network of peers, faculty, or career services programs.<br>Submission tool - front-end: Applicants submit their own materials but they may be aided in this process by peers, faculty, or career services programs. Applicants also indicate their top interests in projects<br>Submission tool - back-end: HF-SRPE program director reviews all applicants prioritized by established funding sources. This provides a second set of eyes on hundreds of applications and reduces some of the burden on mentors hiring for individual projects<br>Hiring documents: Program mentors some autonomy in how they describe project descriptions, set expectations for desired experience, and conducting interviews. HF-SRPE program staff provide resources and accountability to help insure that mentor decisions are aligned with the program's goals and values | QL.1<br>QT.0<br>A.1<br>S.1 | High |
| DL.1 | Expectations of *Division of labor* are seemingly understood by most *community* members but not formally examined | QL.1<br>QT.0 | Med |
|  |  | A.1<br>S.1 |  |
| DL.2 | We routinely receive feedback from the *Community* and believe that there is an adequate *Division of labor* to support the *Mediating artifacts*. | QL.1<br>QT.0<br>A.1<br>S.1 | Med |
| DL.2 | We routinely receive feedback from the *Community* and believe that there is an adequate *Division of labor* to support the *Object*. | QL.1<br>QT.0<br>A.1<br>S.1 | Med |
| DL.3 | The largest change to the *Division of labor* occurred when the program director started doing a preliminary review of applicants. This aided mentors in the early stages of the hiring process and helped relieve some of the burden of reviewing an increasing number of applicants. | QL.1<br>QT.0<br>A.1<br>S.1 | Med-High |
| DL.4 | HF-SRPE likely has similar *Division of labor* compared to other REU programs, especially regarding application materials and hiring documents. However, due to the extraordinarily high number of applications HF-SRPE receives a year, the achievement of *outcomes* such as diversity in hiring may be more sensitive to changes in the *Division of labor*. | QL.0<br>QT.0<br>A.0<br>S.0 | Med |

**Table S4.**
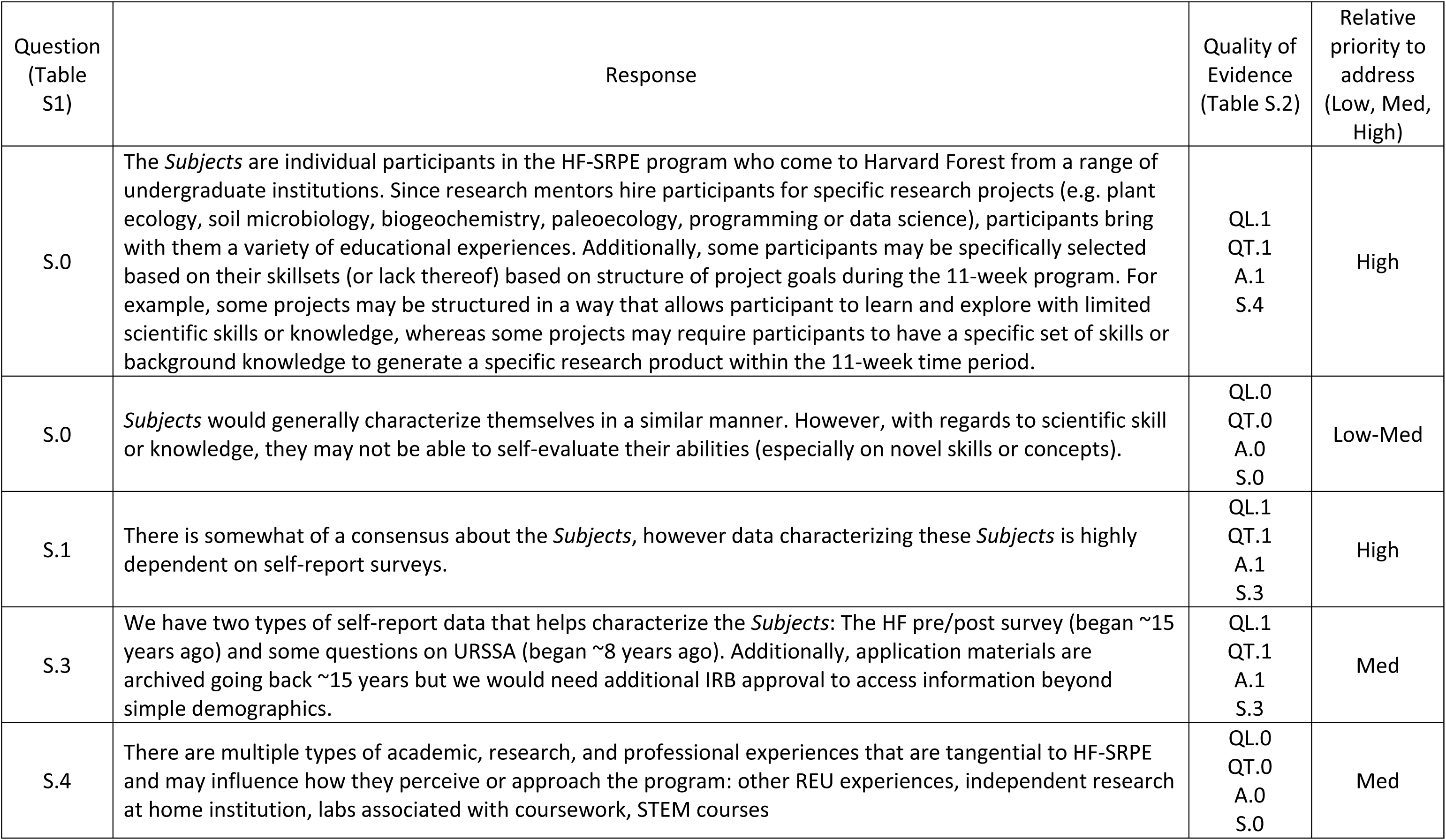
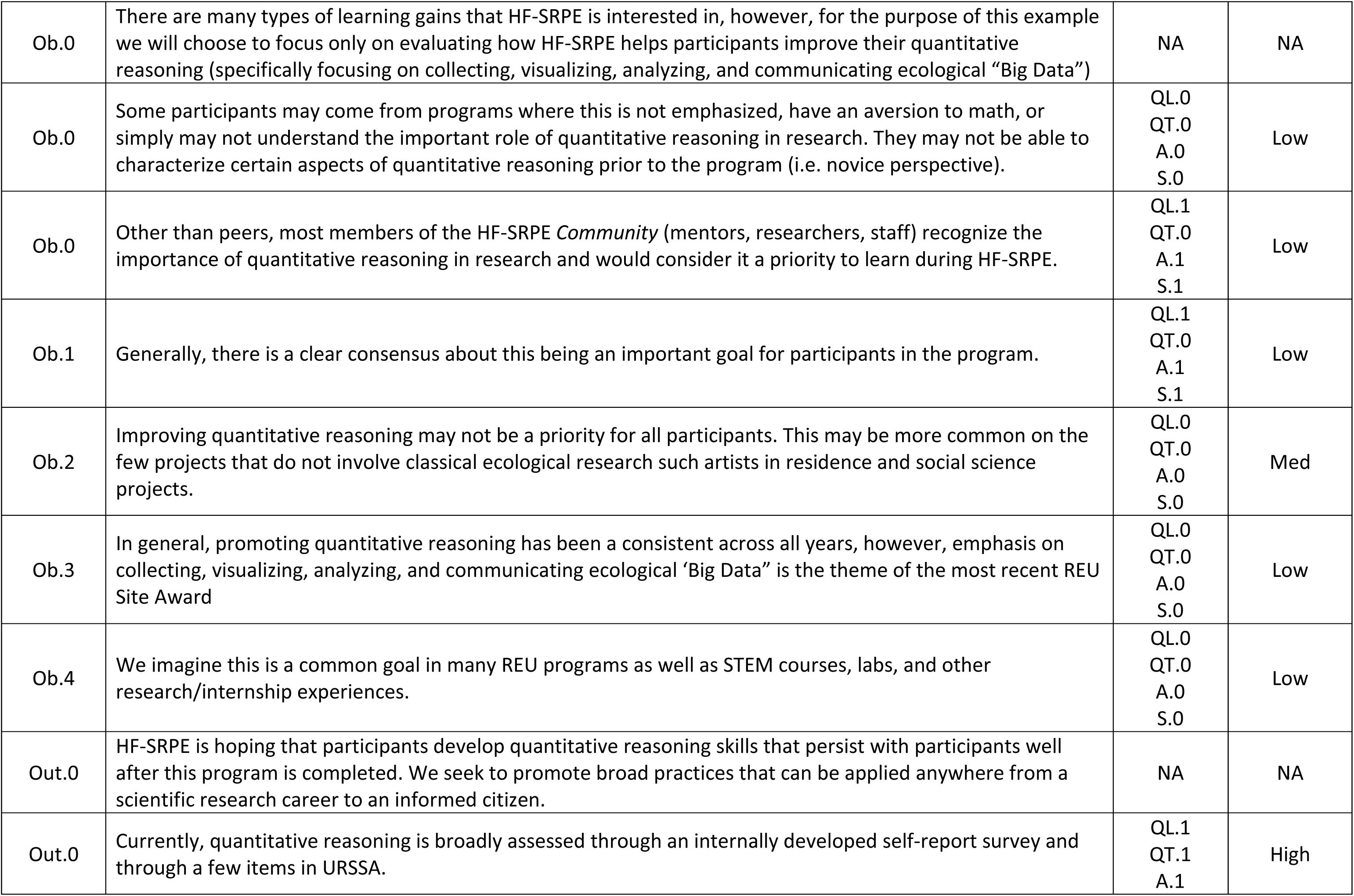
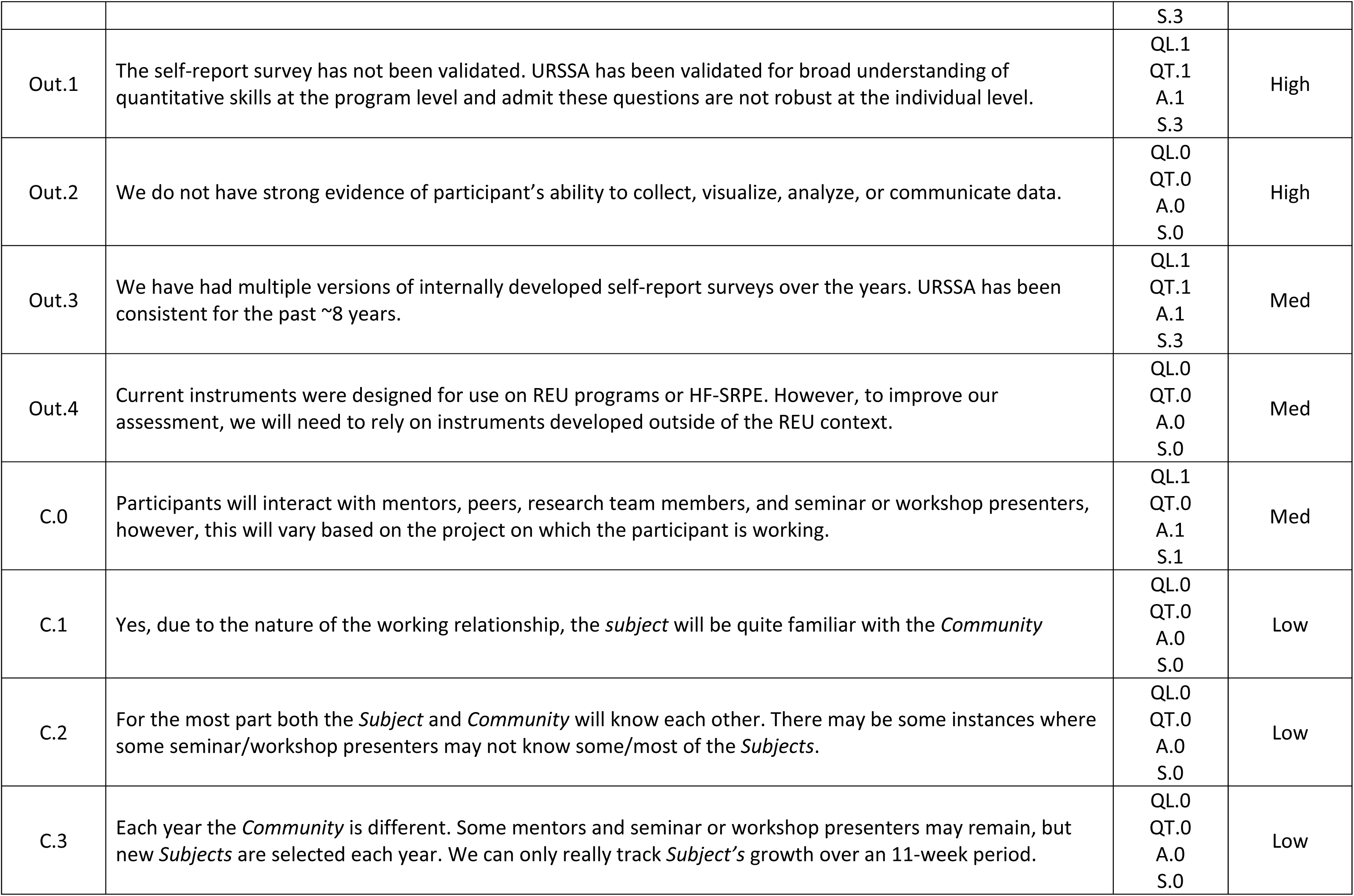
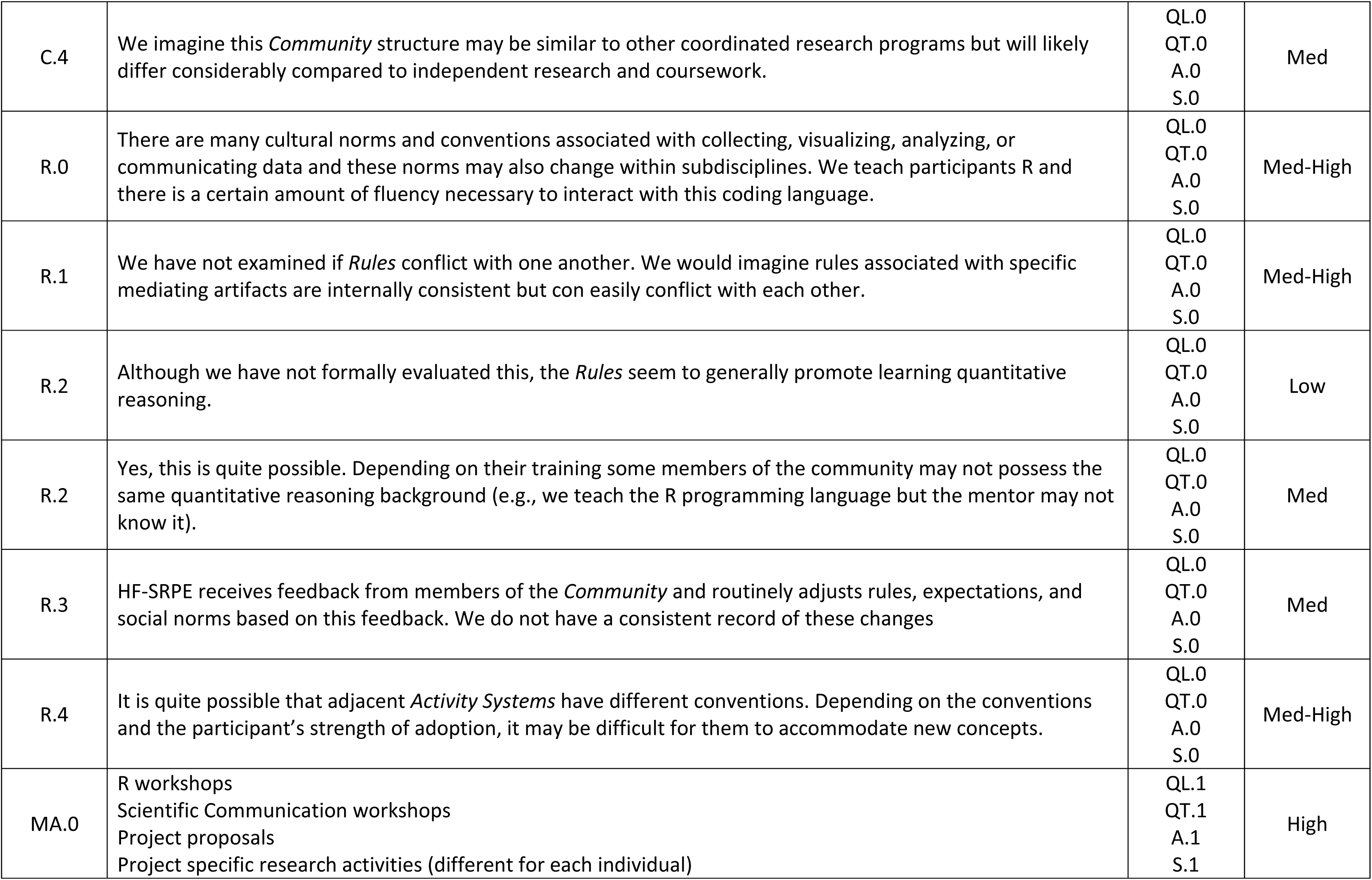
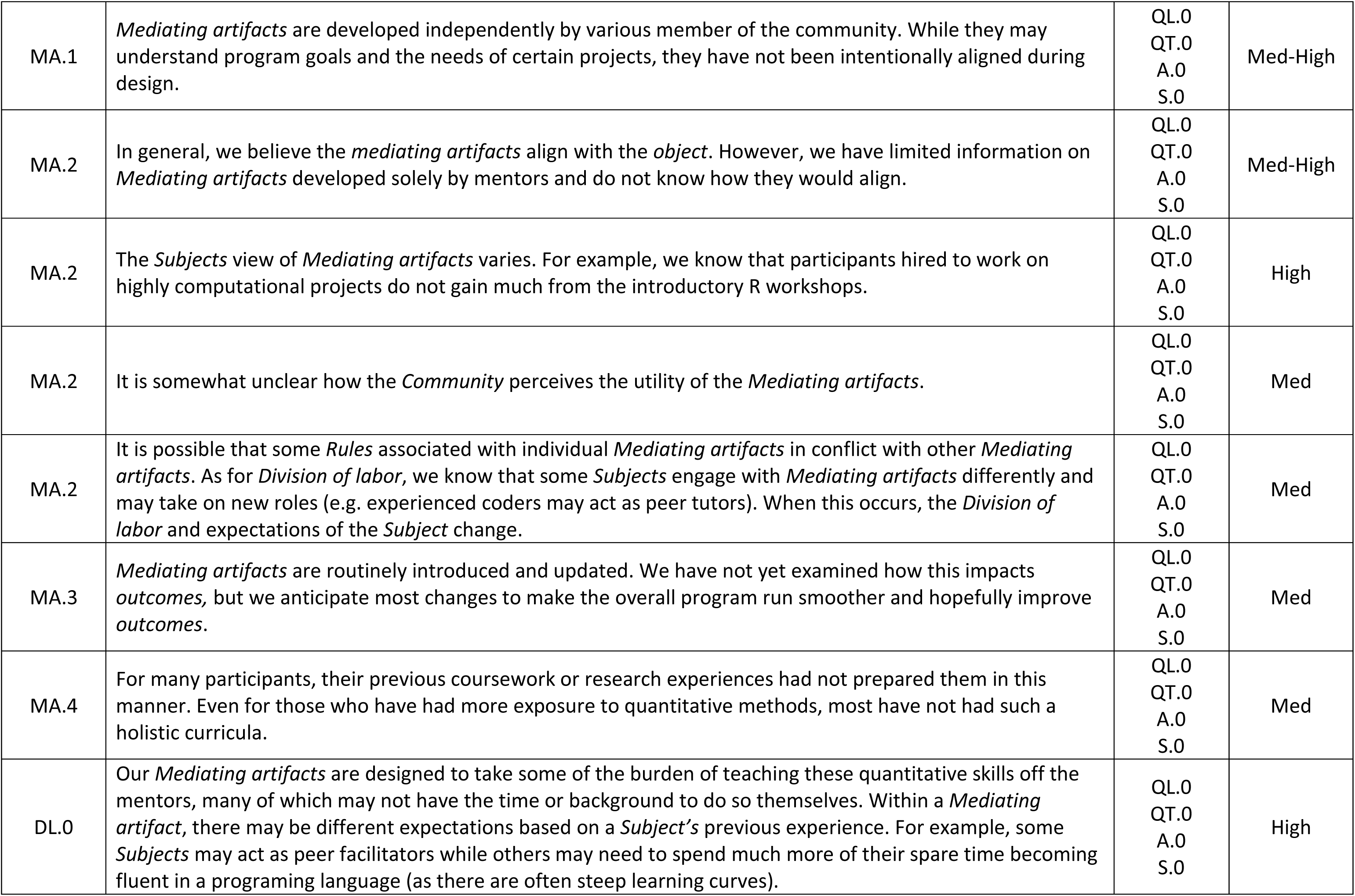
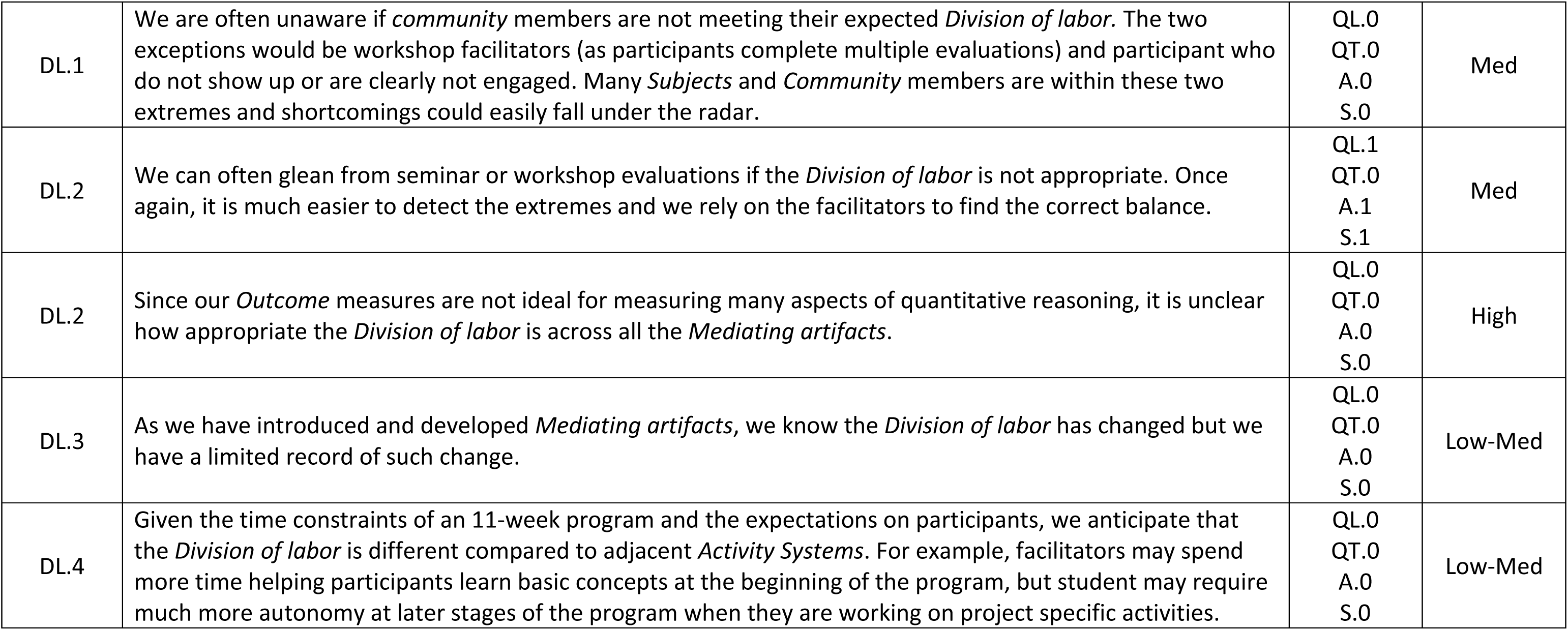
Example responses to a CHAT questionnaire (Table S1) for assessing participant learning gains at HF-SRPE.

**Table S5.** Example responses to the CHAT questionnaire (Table S1) for assessing long-term program impacts of HF-SRPE.

| Question<br>(Table<br>S1) | Response | Quality of<br>Evidence<br>(Table S.2) | Relative<br>priority to<br>address<br>(Low, Med,<br>High) |
| --- | --- | --- | --- |
| S.0 | We tend to attract undergraduate students primarily interested in environmental or ecological careers (as opposed to health professions). Many of our undergraduate students are unclear about their specific career objectives (especially if they are 1<sup>st</sup>- or 2<sup>nd</sup>-year students). For some, this is their first formal research experience and it helps them explore what they like (or dislike) about scientific research. For others with more research experience or firmer career objectives, they take the opportunity to learn skills that will help prepare them for graduate school or the workforce. | QL.1<br>QT.0<br>A.0<br>S.1 | High |
| S.0 | We imagine students would characterize themselves like we have above, however we have not asked them to do so for assessment purposes. | QL.0<br>QT.0<br>A.0<br>S.0 | High |
| S.1 | We are confident in correctly characterizing basic descriptive information (e.g., major, institution type, demographics, etc.), however, we anticipate characterizing career intentions to be difficult. At this point in their undergraduate career, students have many paths to choose from. As they learn about and explore various career options, we imagine that their intents may fluctuate. | QL.0<br>QT.0<br>A.0<br>S.0 | High |
| S.3 | We anticipate that over time, there may be shifts in demands for various skillsets (e.g., working with “Big Data”). As a program, we try to respond to these trends by adjusting projects and programming. It is reasonable to think these changes may lead to a change in how students are characterized. | QL.1<br>QT.0<br>A.0<br>S.1 | Med |
| S.4 | There are multiple types of academic, research, and professional experiences that are tangential to HF-SRPE and may influence how they perceive or approach the program: other REU experiences, independent research at home institution, labs associated with coursework, STEM courses. | QL.0<br>QT.0<br>A.0<br>S.0 | Med |
| Ob.0 | Since 2010, Congress has mandated that we monitor the long-term impact of REU programs. For congress, these goals appear to be focused on building a strong STEM workforce. We generally believe that persistence in STEM is an important, however we also recognize that careers in education/outreach, policy or simply being a more scientifically literate citizen are equally valid long-term goals. | NA | NA |
| Ob.0 | It is somewhat unclear what individual students might state as their long-term goals, but for most, we anticipate that their goals are related to graduate school/employment. However, they may have an alternate perception of success. | QL.1<br>QT.0<br>A.0<br>S.1 | High |
| Ob.0 | We anticipate that research mentors have similar views of long-term success for students. However, as researchers, they may have additional goals such as establishing long-term collaborations with students (either as future graduate students or colleagues). | QL.1<br>QT.0<br>A.0<br>S.1 | Med |
| Ob.1 | While it will depend on the individual student, it is quite possible that there is not a consensus on the long-term goals. It is likely that most of these disagreements will be between the HF-SRPE, student, or mentor and Congressional goals, causing secondary contradictions between the *Object* and *Outcomes* mandated by the America COMPETES Act of 2010. | QL.0<br>QT.0<br>A.0<br>S.0 | High |
| Ob.2 | Since students apply to take part in HF-SRPE, we'd expect that most of their long-term goals are in alignment with the program. However, tensions could arise during *Mediating artifacts* if the intent was not aligned with the *Subject's* perception of long-term goals (i.e. a student with no interest in graduate were required to write or participate in a workshop about graduate school admission essays). | QL.0<br>QT.0<br>A.0<br>S.0 | Low |
| Ob.3 | We do not anticipate large cultural shifts in long-term goals. | QL.0<br>QT.0<br>A.0<br>S.0 | Low |
| Ob.4 | Similar goals are common throughout U.S. culture; however, students may be a part of a community or home institution that may hold a different set of values. | QL.0<br>QT.0<br>A.0<br>S.0 | Low |
| Out.0 | Success may mean multiple things to multiple people. Just because long-term success is not achieved under one metric, does not mean that a student did not have a successful experience. Additionally, there are many other factors that contribute to the "long-term success" of students and it would be presumptuous to think that an 11-week program is the only factor leading to this metric. Additionally, people have different paths to success and may appear more or less successful depending on when the measurements are taken. | NA | NA |
| Out.0 | Our assessment priority for long-term success will be dictated by the America COMPETES Act of 2010 which requires the tracking of students for STEM matriculation and employment for at least three years following graduation. We have an annual survey that is sent to all program alumnae/i (since 2001) that tracks enrollment and attainment of degrees, and STEM employment. | NA | NA |
| Out.1 | This *Outcome* is a requirement of our funding agency and HF-SRPE has the discretion to choose how we measure the *Outcome*. Our current measurement relies on self-report data. We have a convenience sample which decreases over time as we loose track of former students (e.g., out of date contact information, name changes) | QL.1<br>QT.1<br>A.1<br>S.1 | High |
| Out.2 | Our *Outcome* measures only have face validity. | QL.0<br>QT.0<br>A.1<br>S.0 | Med |
| Out.3 | Our alumni survey has remined relatively consistent since 2010. | QL.1<br>QT.1<br>A.1<br>S.0 | Low |
| Out.4 | We developed this survey in-house but have been exploring other techniques to track these *Outcomes*. One promising avenue for improvement is to transition to techniques used by college alumni associations. | QL.0<br>QT.0<br>A.0<br>S.0 | Med |
| C.0 | Students will interact with mentors, peers, research team members, and seminar or workshop presenters, however, this will vary based on the project on which the student is working. | QL.1<br>QT.0<br>A.1<br>S.1 | Med |
| C.1 | Yes, because of the nature of the working relationship, the *Subject* should be quite familiar with the *Community* | QL.0<br>QT.0<br>A.0<br>S.0 | Low |
| C.2 | For the most part both the *Subject* and *Community* will know each other. There may be some instances where some seminar or workshop presenters may not know some or most of the *Subjects*. | QL.0<br>QT.0<br>A.0<br>S.0 | Low |
| C.3 | Each year the *Community* is different. Some mentors and seminar or workshop presenters may remain, but new *Subjects* are selected each year. We can only really track each *Subject's* growth over an 11-week period. | QL.1<br>QT.0<br>A.1<br>S.1 | Low |
| C.4 | We imagine this *Community* structure may be like that of other coordinated research programs but will likely differ considerably compared to independent research and coursework. | QL.0<br>QT.0<br>A.0<br>S.0 | Med |
| R.0 | Our goal is to treat students as employees and colleagues rather than “undergraduate students”. The aim is to model professional behavior and provide support to students (some of which this is their first job) so that they eventually feel a sense of autonomy and accountability for their actions. | QL.1<br>QT.0<br>A.1<br>S.1 | Med-High |
| R.1 | Since students are living at the Harvard Forest, there is sometimes difficulty delineating work from recreation. Social norms are established between housemates, peers, and supervisors which are unique to each cohort. This is challenging to navigate, and it is very easy for individuals to receive social signals that are at odds with the intent of the program. | QL.1<br>QT.0<br>A.1<br>S.1 | Med-High |
| R.2 | Although we try to establish consistent rules and expectations, there are instances where rules and norms may contradict each other. This often happens in response to an incident at work or in the residences that require enforcement of rules and policies. In these instances, the actions of a few individuals may cause the group to feel a loss of autonomy. | QL.1<br>QT.0<br>A.1<br>S.1 | Low |
| R.2 | No, students bring with them their own set of values which could be at odds with the norms that HF-SRPE is trying to establish. We try to mitigate these conflicts by being transparent about the rationale for certain expectations and open to dialogue (although this is easier said than done). | QL.0<br>QT.0<br>A.0<br>S.0 | Med |
| R.3 | Although the intent is to be consistent with rules, expectations, or cultural norms from year to year (modeled after the Harvard Forest research community), there is inherently some variability. | QL.1<br>QT.0<br>A.1<br>S.1 | Med |
| R.4 | HF-SRPE has a similar feel to academic research cultures on traditional campuses, however, there are many aspects related to a rural biological field station that creates a distinct set of rules and expectations. For example, feelings of isolation and irritability (i.e., cabin fever) are common among students and we try to be cognizant about how we provide support to students as they transition to the new environment (and the norms associated with them). | QL.0<br>QT.0<br>A.0<br>S.0 | Med-High |
| MA.0 | We provide several structured and informal opportunities for students with regards to graduate school and career opportunities.<br>• Career panel<br>• Networking with researchers | QL.1<br>QT.0<br>A.1<br>S.3 | High |
|  | Other structured activities focus on broader skills useful for scientific careers<br>• Independent research project (with a research mentor)<br>• Project proposals<br>• Science Communication workshop<br>• Blogging<br>• Poster workshop<br>• R/programming workshop<br>• Research seminars<br>Additionally, there are many more “one-off” opportunities that are created by research mentors or based on student interest. These are often in response to an individual’s summer/career goals. |  |  |
| MA.1 | When designing the programming, we consider how, and professional development activities support students with respect to the mission of the program and student long-term goals. | QL.1<br>QT.1<br>A.1<br>S.2 | Med-High |
| MA.2 | We continuously solicit feedback from students and mentors to monitor how students’ short term and long-term goals are being supported. | QL.1<br>QT.1<br>A.1<br>S.2 | Med-High |
| MA.2 | Students perception of utility often depends on their skillsets or experiences. When designing an activity, we try to accommodate a range of skill levels. | QL.1<br>QT.0<br>A.1<br>S.1 | High |
| MA.2 | The *Community* generally finds these professional development activities useful. However, individual research mentors may prioritize research above some required activities. | QL.0<br>QT.0<br>A.0<br>S.0 | Med |
| MA.2 | In general, we believe that our *Rules* align these *Mediating artifacts* and how they are accomplished. | QL.0<br>QT.0<br>A.0<br>S.0 | Med |
| MA.3 | Based on feedback, we continuously introduce, remove, and revise *Mediating artifacts*. It is unclear how these changes are related to long-term success. | QL.0<br>QT.0 | Med |
|  |  | A.0<br>S.0 |  |
| MA.4 | These *Mediating artifacts* are common to other research programs and educational settings. We find that even if a student has completed a similar activity before, the repetition is useful as they may gain a different perspective the second (third or more) time around. | QL.0<br>QT.0<br>A.0<br>S.0 | Low |
| DL.0 | The research mentor and student are asked to outline expectations for the project proposal at the beginning of the summer. *Division of labor* is somewhat variable among research mentors which is why HF-SRPE provides formal programming to help ensure a consistent exposure to professional development resources. Sometimes, a student will maintain a research relationship with their mentor after the end of the summer (often resulting in a research project such as a poster, undergraduate thesis, or manuscript).<br>For other professional development activities, HF-SRPE strives to provide resources for students that their research mentors may not otherwise have the time or expertise to provide. | QL.1<br>QT.0<br>A.1<br>S.1 | Med |
| DL.1 | Although we have a proposal that outlines expectation for each student's project, we do not revisit these documents to evaluate if the agreed upon *Division of labor* was met. Our reluctance to analyze these documents is due to the how these documents are formatted (some projects require a lot of structure while others are more trial-and-error) and that research goals may change rapidly throughout the summer. | QL.0<br>QT.0<br>A.0<br>S.0 | High |
| DL.2 | Most of the time, students and mentors find the *Division of labor* is appropriate. However, we do have mentors and students come to program staff when they feel that expectations are not being met. Program staff act as mediators to resolve any conflicts and provide alternative to help both side move forward in a productive manner. | QL.0<br>QT.0<br>A.0<br>S.0 | Med |
| DL.2 | Currently it appears that the *Division of labor* throughout the program is appropriate. | QL.1<br>QT.0<br>A.1<br>S.1 | High |
| DL.3 | Project proposals were introduced in response to student feedback that the *Division of labor* was not being met by some research mentors. The introduction of this *Mediating artifact* helped to clarify expectations. | QL.1<br>QT.0<br>A.1<br>S.1 | Low-Med |
| DL.4 | For both students and research mentors, the *Division of labor* may be different than what they are used to in other settings. We stress that HF-SRPE students have intellectual involvement in the project and should not be | QL.0<br>QT.0<br>A.0 | Med-High |
|  | viewed as grunt labor. This may cause a shift in how some individuals view their role and we try to provide support to aid in such a transition. | S.0 |  |

